# The effect of experimental lead pollution on DNA methylation in a wild bird population

**DOI:** 10.1101/851998

**Authors:** Hannu Mäkinen, Kees van Oers, Tapio Eeva, Veronika N. Laine, Suvi Ruuskanen

## Abstract

Anthropogenic pollution is known to negatively influence an organism’s physiology, behavior and fitness. Epigenetic regulation, such as DNA methylation, has been hypothesized as one mechanism to mediate such effects, yet studies in wild species are lacking. We first investigated the effects of early-life exposure to the heavy metal lead (Pb) on DNA methylation levels in a wild population of great tits (*Parus major*), by experimentally exposing nestlings to lead at environmentally relevant levels. Secondly, we studied the effects of heavy metal exposure in a population close to a copper smelter, where birds suffer from pollution-related decrease in food quality. For both comparisons, the analysis of about million CpGs covering most of the annotated genes, revealed that regions enriched for developmental processes showed pollution-related changes in DNA methylation, but the results were not consistent with binomial and beta binomial regression. Our study indicates that post-natal anthropogenic heavy metal exposure can affect methylation levels of development related genes in a wild bird population.

## Introduction

Epigenetic control of gene expression, such as DNA methylation, is increasingly recognized as playing a major role in many different cellular processes. DNA methylation is the addition of a methyl (-CH_3_) group to the 5’ carbon site of cytosines catalyzed by DNA-methyltransferases that occurs mainly at CpG sites in animals. Especially in CpG islands within promotor regions, DNA methylation is found to be negatively associated with gene expression. Epigenetic changes are linked to variation in phenotype and behavior and are associated with prevalence for various diseases (Angers et al. 2010, Rosenfeld 2010, Skinner et al. 2010).

Methylation patterns can be affected by various environmental factors such as maternal nutrition and maternal care (e.g. Weaver et al. 2004, Heijmans et al. 2008, Faulk and Dolinoy 2011, Feil and Fraga 2012), but also by various pollutants and other early-life stressors, both pre- and postnatally, as discovered in humans and mouse models (reviewed by Cheng et al. 2012, Head et al. 2012, Head 2014, Ray et al. 2014, Ruiz-Hernandez et al. 2015). The potential effects of environmental factors on epigenetic regulation is highly important for ecological and ecotoxicological fields, but research in wild vertebrate populations is only emerging (Bossdorf et al. 2008, Head et al. 2012, Liebl et al. 2013, Wenzel and Piertney 2014, Riyahi et al. 2015, Rubenstein et al. 2016, Verhoeven et al. 2016). However, epigenetics can significantly improve our understanding of the mechanisms underlying natural phenotypic variation and the responses of organisms to environmental change (Verhoeven et al. 2016). Furthermore, potential transgenerational epigenetic effects could explain why populations are slow to recover even after pollution removal (Head 2014). A major barrier for genome-wide methylation studies has been the lack of adequate genomic resources such as sequenced genomes in non-model species (Viitaniemi et al. 2019).

Heavy metals, such as lead (Pb), are global, persistent human-induced pollutants that are among the potential contaminants affecting DNA methylation status (reviewed in Bihaqi 2019). For example in human epidemiological studies, developing fetuses show a decrease in global methylation levels as a result of historical maternal lead exposure and accumulation (Pilsner et al. 2009, Wright et al. 2010). Furthermore, in rat, mouse and monkey models, experimental peri- and post-natal lead exposure decreases DNA methyltransferase activity and affects DNA methylation, which are subsequently related to behavioral alternations (Wu et al. 2008, Faulk et al. 2013, Faulk et al. 2014, Luo et al. 2014, Sanchez-Martin et al. 2015, Singh et al. 2018, Nakayama et al. 2019). In birds, metal exposure has been found to affect offspring growth (Burger and Gochfeld 2000) and multiple aspects of physiology, including stress hormone and stress protein levels (Eeva et al. 2014), oxidative stress levels (Koivula and Eeva 2010) and immune function (reviewed in Boyd 2010). However the potential epigenetic alterations by early-life exposure to metal pollution, potentially underlying such effects in birds, have not been studied.

In addition to the direct effect of metals, large-scale metal pollution can decrease resource availability and quality in wild populations (Eeva and Lehikoinen 2004, Eeva et al. 2005), which could subsequently also influence methylation patterns: Along with toxicants, altered nutrition and diet, especially diet poor in methyl donors, are well-known epigenetic modifiers in animal models (reviewed in Choi and Friso 2010, Rosenfeld 2010, Konycheva et al. 2011). Also protein or lipid-altered diets can cause major changes in the epigenome (Burdge et al. 2007, Aagaard-Tillery et al. 2008, Choi and Friso 2010). All in all, we expect populations inhabiting polluted environments to have altered DNA methylation patterns, either due to direct or indirect pollution effects.

Here we investigated whether experimental and anthropogenic early-life exposure to the heavy metal lead (Pb) alters genome-wide DNA methylation status in a wild population of great tits (*Parus major*). More specifically, we experimentally exposed nestlings to dietary lead at levels found close to active pollution sources in Europe and compared to respective controls. The exposure covered the whole pre-fledging period. Secondly, we studied methylation patterns using nestlings from a population close to an anthropogenic pollution source, copper smelter (Eeva et al. 1997). Around the smelter, nestlings are exposed to multiple metals (in low concentrations) and experience an altered nutritional quality and quantity compared to controls. In the latter case we therefore studied the combined effect of altered nutrition (indirect effect of metal pollution) and direct, low, metal exposure on DNA methylation. In our recent work using the same experimental protocol we found that lead exposure and altered nutrition during nestling development lead to changes in e.g. growth, oxidative stress markers, stress protein levels and vitamin metabolism, but the mechanisms, potentially epigenetic regulation are not understood (Eeva et al. 2014, Rainio et al. 2015b, Ruiz et al. 2016). We used reduced representation bisulfite sequencing (RRBS) to characterize methylation levels, and the sequenced genome of the study species (Derks et al. 2016, Laine et al. 2016, Verhulst et al. 2016) to assess methylation differences in different gene features and their potential functional significance. Given that there are not yet established golden standards for analyzing methylation data in an ecological context, we used and compared two frequently used analytical tools to detect differentially methylated sites (Wreczycka et al. 2017). To avoid sacrificing the individuals, we used blood as a source tissue; previous studies suggest that blood shows similar methylation patterns as brain tissue in the study species (e.g. 80% similarity between brain and blood methylation in CpGs Derks et al. 2016, Verhulst et al. 2016).

## Methods

### Study species

The great tit (*Parus major)* is a small passerine bird and a model species in ecological and evolutionary research, with ample ecological and genetic background information available. It is an insectivorous non-migratory bird that commonly breeds in nest boxes, making it an ideal species for experimental manipulations. Importantly, as one of the only non-domesticated bird species, both the genome and methylome are available (Bouwhuis et al. 2010, Derks et al. 2016, Laine et al. 2016, Verhulst et al. 2016).

### Lead treatment protocol and sampling

The lead exposure, dosages and sampling are described in detail in (Eeva et al. 2014, Rainio et al. 2015b, Ruuskanen et al. 2015). Briefly, breeding was monitored to record hatching dates of great tit chicks in a population with low pollution levels in southwestern Finland (Turku, 60°26’N, 22°10’E). From day 3 after hatching (hatch date = 0) until day 14 (i.e. in total 12 days) whole broods were subjected to lead with daily oral dosing with the following treatments: HIGH dose (4 µg Pb/g body mass, N = 15 broods) or CONTROL (distilled water, N = 15 broods). The dose represented environmentally relevant exposure levels occurring in polluted areas in Europe. The exposure period covered most of the post-hatching nestling period, i.e. most important developmental period in altricial birds. The third group (hereafter SMELTER) consisted of nests (N = 19 broods) in the vicinity (<2 km) of Harjavalta copper smelter (61°20’N, 22°10’E), where there is a long-term exposure of several metals (e.g., lead, arsenic, cadmium, copper, nickel) and generally lower food availability and quality (Eeva and Lehikoinen 2004). These nests were monitored in the same way as described above and nestlings were dosed with distilled water.

Blood samples were collected from 7-day old nestlings for sex-determination (following Griffiths et al. 1998) and measurements were taken of multiple physiological indices (Eeva et al. 2014, Rainio et al. 2015a). Fresh fecal samples were collected for measuring metal concentrations (see below). Whole blood samples were collected directly in liquid nitrogen from nestlings at age of 14 days (i.e. after 12 days of treatment) for analyses of DNA methylation status and physiological indices. Samples were stored at −80°C until analysis. Ten unrelated samples from female nestlings in HIGH and CONTROL groups, and eight samples from SMELTER group were selected randomly (total N = 28 samples). The experiment was conducted under licenses from the Animal Experiment Committee of the State Provincial Office of Southern Finland (license number ESAVI/846/ 04.10.03/2011) and the Centre for Economic Development, Transport and the Environment, ELY Centre Southwest Finland (license number VARELY/149/07.01/2011).

### Metal analyses

For detailed analyses, see Eeva et al. (2014). Briefly, two fecal samples (one male and one female) from the same brood were combined to assess brood level metal exposure (total N = 35 broods). The determination of metal concentrations (As, Cd, Cu, Ni, Pb) was conducted with ICP-MS with detection limit of 1 ppt (ng/l) and below. The calibration of the instrument was done with a commercial multi-standard from Ultra Scientific, IMS–102, ICP-MS calibration standard 2 and certified reference materials were used for method validation. Data was analyzed with GLMs (SAS 9.4) with Tukey post-hoc tests.

### DNA isolation

DNA isolation was performed at the Center of Evolutionary Applications (University of Turku, Finland). We used whole blood samples, which can be acquired without sacrificing the individuals. In birds erythrocytes have nuclei and therefore >95% of the gained DNA is from erythrocytes. DNA was extracted from c. 10 − 20 μl whole blood using the salt extraction method modified from Aljnabi & Martinez (1997). Extracted DNA was treated with RNase-I according to the manufacturer’s protocol. DNA concentrations were measured fluorometrically with a Qubit High Sensitivity kit (ThermoFisher Scientific) and we assessed DNA integrity by running each DNA sample on an agarose gel.

### RRBS library preparation

We used a reduced representation bisulfite sequencing (RRBS) approach, which enriches the regions of the genome that have a high <u>CpG</u> content (Meissner et al. 2005). It was previously shown in the study species that the vast majority of the methylated cytocines (97%) were derived from CpG sites in blood (Derks et al. 2016). Sequencing was conducted at the Finnish Microarray and Sequencing Center in Turku, Finland. The library preparation was started from 200 ng of genomic DNA and was carried out according to a protocol adapted from Boyle et al. (2012). The first step in the workflow involved the fragmentation of genomic DNA with MspI where the cutting pattern of the enzyme (C^CGG) was used to systematically digest DNA to enrich for CpG dinucleotides. After a fragmentation step, a single reaction was carried out to end repair and A-tail (required for the adapter ligation) the MspI digested fragments using Klenow fragment (3’ => 5’ exo), following the purification of A-tailed DNA with bead SPRI clean-up method (AMPure magnetic beads). A unique Illumina TruSeq indexing adapter was then ligated to each sample during adapter ligation step to be able to identify pooled samples of one flow cell lane. To reduce the occurrence of adapter dimers, a lower concentration of adapters (1:10 dilution) was used than recommended by the manufacturer. These ligated DNA fragments were purified with the bead SPRI clean- up method before putting samples through bisulfite conversion to achieve C-to-U conversion of unmethylated cytosines, whereas methylated cytosines remain intact. Bisulfite conversion and sample purification were done according to the Invitrogen MethylCode Bisulfite Conversion Kit. Aliquots of converted DNA were amplified by PCR (16 cycles) with Taq/Pfu Turbo Cx Polymerase, a proofreading PCR enzyme that does not stall when it encounters uracil, the product of the bisulfite reaction, in the template. PCR-amplified RRBS libraries were purified using two subsequent rounds of SPRI bead clean-ups to minimize primer dimers in the final libraries. The high quality of the libraries was confirmed with Advanced Analytical Fragment Analyzer and the concentrations of the libraries were quantified with Qubit® Fluorometric Quantitation, Life Technologies. We selected fragment sizes ranging between 150 − 1000 bp (average sizes were 250-350 bp) for sequencing.

### Sequencing

The samples were normalized and pooled for the automated cluster preparation, which was carried out with an Illumina cBot station. The 28 libraries were randomly combined in three pools, 10 or 8 samples in each pool and sequenced in 3 lanes. The samples were sequenced with an Illumina HiSeq 2500 instrument using TruSeq v3 sequencing chemistry. Paired-end sequencing with 2 x 100 bp read length was used with 6 bp index run.

### Bisulfite sequencing analysis

The initial quality check with Fastqc (Andrews, 2010) indicated presence of Illumina universal adapter contamination and low quality (Q < 20) bases in the 3’ end of the raw reads. The adapter sequences were removed with Cutadapt (Martin 2014) and the low quality bases were filtered using Condentri (Smeds and Kunstner 2011) with default settings. The quality filtered reads were then mapped against the Great tit reference genome (Assembly Parus_major1.0.3; NCBI Bioproject PRJNA208335, Laine et al. 2016) using Bismark aligner with default parameters (L, 0, -0.2) allowing 2−3 mismatches or a comparable number of indels per 100 bp read (Krueger and Andrews 2011). Methylation information was extracted from alignment files using the bismark_methylation_extractor tool (Krueger and Andrews 2011). The resulting methylation levels per base pair were inspected to detect potential methylation bias in the beginning and in the end of read 1 and 2 (Hansen et al. 2012). There was lower methylation in the beginning and higher in the end of read 2. Therefore, the first four bases and the last base were removed from the read 2 for subsequent analyses (supplementary figure 1). On average, we recovered 16.08 million raw reads (range 13.55-21.15) from each RRBS library and after quality filtering 11.72 million reads remained (range 10.12-15.06). On average 6.36 million (54%) of the quality filtered reads were uniquely mapped against the Great tit reference genome. This translates to an average 322.45 million cytosine bases analyzed, of which 179.95 million cytocines (56%) were methylated (Supplementary table 1). We estimated the bisulfite conversion rate by aligning the reads against great tit mitochondrial DNA, which is mostly un-methylated (Mechta et al. 2017) and calculated the conversion rate as 1-methylation% in CpG context. The bisulfite conversion rate was 97.7-99.1%.

**Figure 1.**
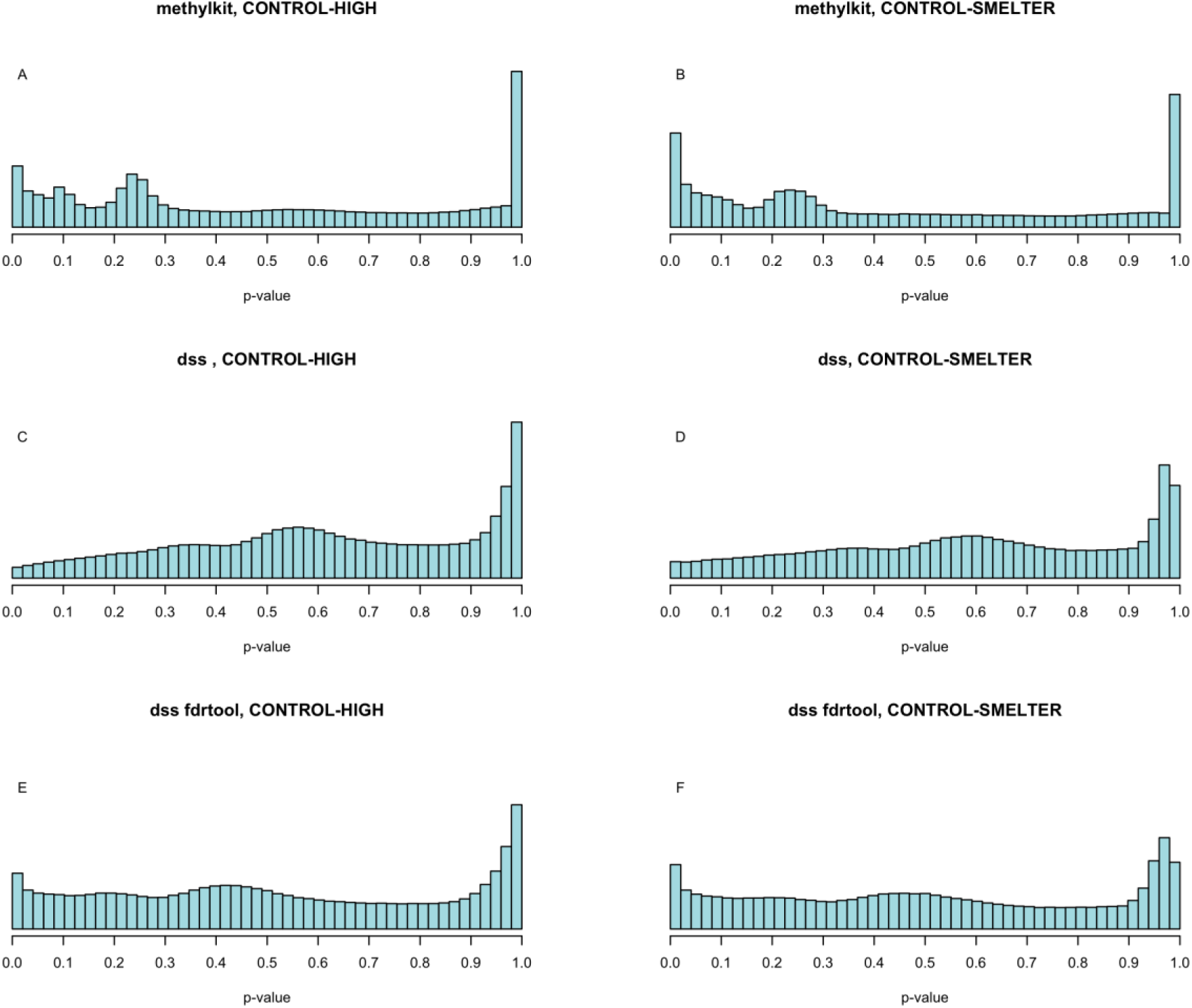
P-value histograms of the binomial (methylkit, A, B) and beta-binomial regression (dss, C, D) in CONTROL-HIGH and CONTROL-SMELTER comparisons. The p-values of the beta binomial regression were re-calculated based on the test statistics as implemented in the fdrtool R-package (E, F).

**Table 1.**
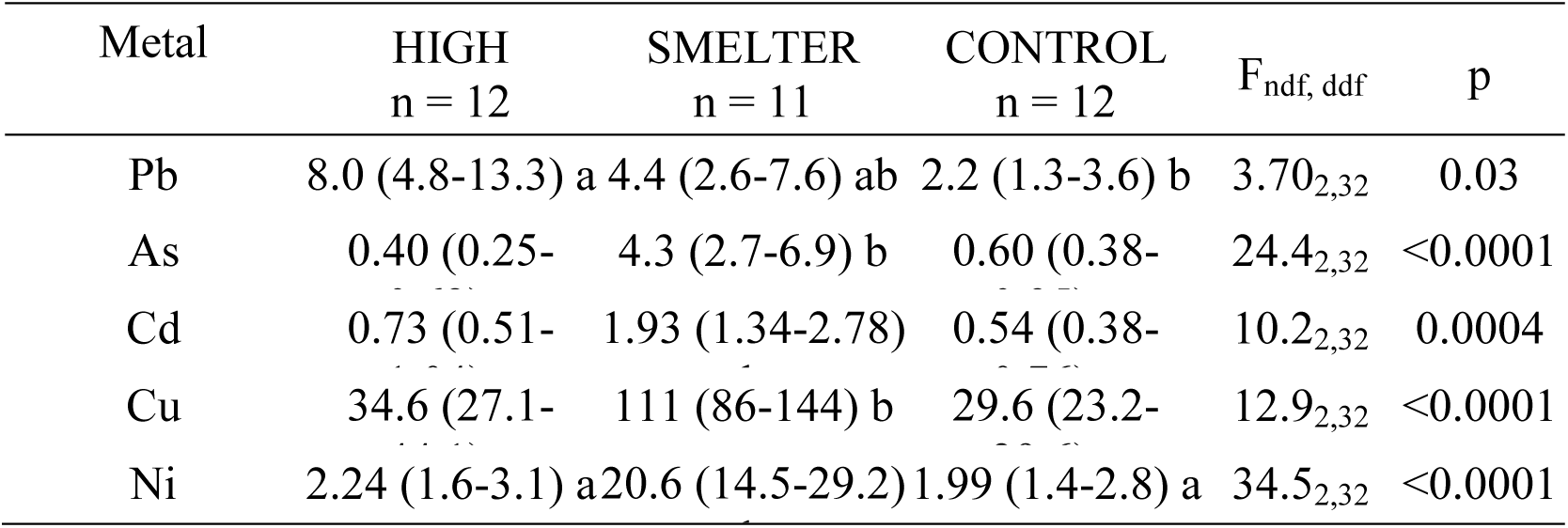
Metal concentrations (µg/g, dry weight) in feces of seven day old *Parus major* nestlings in the three treatment groups. The values are geometric means with 95% CIs. GLM and Tukey’s test: means with the same letter are not significantly different. N indicates number of broods.

In order to call methylated CpG sites from the Bismark methylation extractor files, the function *readBismarkCoverage* in the R package Methylkit (Akalin et al. 2012) was used. Using a minimum coverage threshold of 10, on average 1309860 (range 1062814-1545820) methylated CpG sites were obtained for the CONTROL-HIGH comparison and 1344110 (range 1062814-1774992) CpG sites for the CONTROL-SMELTER comparison. The CpGs were then filtered by extreme coverage to remove e.g. potential PCR duplicates using 99.9% percentile upper threshold as implemented in the function *filterByCoverage* in R package Methylkit. Methylated CpG sites were also median normalized to take into account differing library sizes using *normalizeCoverage* function in R package Methylkit. Finally, CpG sites were united such that the final data set contained only CpG sites covered by a minimum of seven individuals per group. The final data sets comprised 1023725 and 903449 CpG sites for CONTROL-HIGH and CONTROL-SMELTER comparisons, respectively. 879056 CpG sites were shared between these two comparisons.

Two methods were used for the identification of differentially methylated CpG sites. First, a generalized linear model was used as implemented in the R package Methylkit. This method assumes that the methylated and un-methylated counts follow a binomial distribution and the effect of group/treatment can be estimated with a log-likelihood test (Akalin et al. 2012). Second, a generalized linear model assuming beta binomial distribution was used taking into account potential overdispersion by estimating a gene-specific shrinkage operator as implemented in R package dss (Wu et al. 2013; Feng et al. 2014). In both methods, the model was fitted for each CpG site separately and we compared the pairwise methylation differences between the CONTROL and HIGH and CONTROL and SMELTER groups. The model fits were evaluated by inspecting the resulting p-value histograms. Under a proper null model one would expect that the p-value histogram follows a uniform distribution, but if there is an effect of e.g. treatment then a surplus of small p-values is expected (Fodor et al. 2007, Barton et al. 2013, Garamszegi and de Villemereuil 2017). Deviations from the uniform distribution may provide information about the misspecification of the model or problems in the data. Differentially methylated regions (DMRs) or clusters of differentially methylated CpG sites were identified based on the results of both binomial and beta binomial models. The following criteria were used to identify DMRs: (i) minimum size of a DMR = 50 bp, (ii) minimum number of CpGs in the DMR = 3 and (iii) the percentage of CpGs with p-value < 0.01 in the cluster = 50%. The identification of DMRs was conducted in the R package dss.

#### Annotation of differentially methylated regions

The location and association of the CpGs with a given genomic feature was determined using the Great tit genome assembly and annotation 1.1 (Laine et al. 2016). More specifically, each CpG was annotated with respect to location in genes, exons and in coding sequence (CDS). The annotation of the DMRs was conducted using the *IntersectBed* options in the BedTools package to identify the overlapping genomic features (Quinlan and Hall 2010). STRING database (Szklarczyk et al. 2015) was used to identify gene ontology categories associated with the DMRs. A hierarchical clustering with a user-specified cutoff value C (0.5) was used as implemented in REVIGO database for merging of semantically similar GO categories corresponding to 1% chance of merging two randomly generated categories (Supek et al. 2011). The lowest p-values of the initial enrichment analyses were used to select a representative GO term for each merged category.

## Results

### Metal exposure

Our dietary lead treatment (HIGH lead) significantly increased fecal lead concentrations compared to the CONTROL (Table 1). At the SMELTER site, we found intermediate fecal lead levels, not significantly different from either HIGH or CONTROL (Table 1). In the SMELTER area, concentrations of other measured heavy metals (As, Cu, Cd, Ni) were higher than in CONTROL or HIGH treatment (Table 1).

### DNA methylation

#### Overall methylation differences among the treatment groups

The range of mean and median coverage was 25.94-33.31 and 17-27 in the CONTROL-HIGH comparison. The respective statistics were 33.73-41.50 and 27-37 in the CONTROL-SMELTER comparison. The average methylation percentages across all CpGs were 27.97 and 28.01 in CONTROL-HIGH lead comparison, and 26.82 and 26.67 in CONTROL-SMELTER comparison. The mean difference in methylation in CONTROL-HIGH and CONTROL-SMELTER comparisons were −0.0013% and 0.0015%, respectively. There were no marked differences in the methylation percentage using 2% cutoff between the major chromosomes in either comparison (Supplementary figure 2). Also, there were no clear patterns in sample clustering in either of the comparisons based on hierarchical clustering or principal components analysis (Supplementary figure 3, Supplementary figure 4).

**Figure 2.**
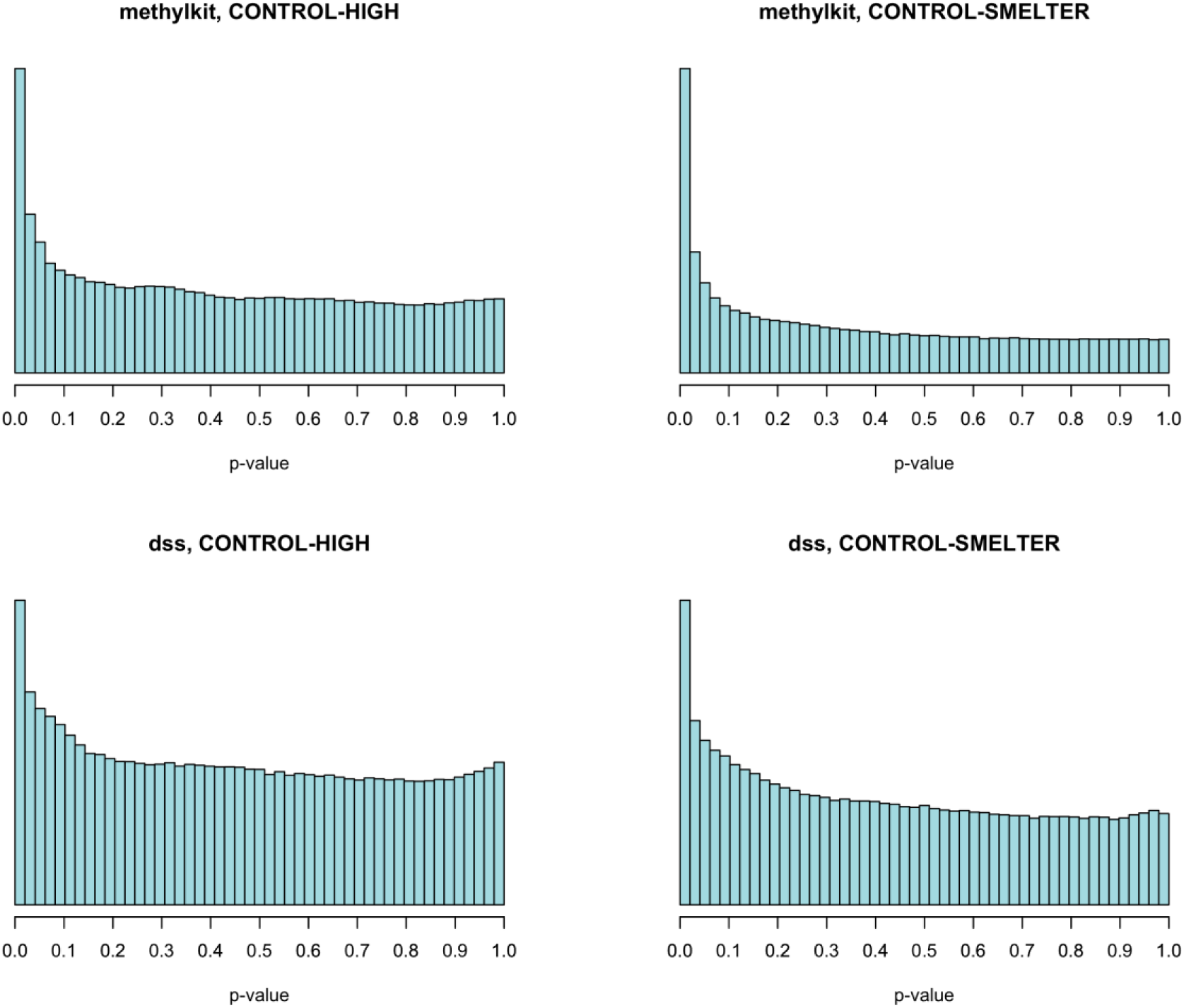
The p-value histograms after low coverage filtering in binomial regression (methylkit) and in beta binomial regression (dss). The p-values in beta binomial regression are based on the re-calculated p-values in fdrtool.

**Figure 3.**
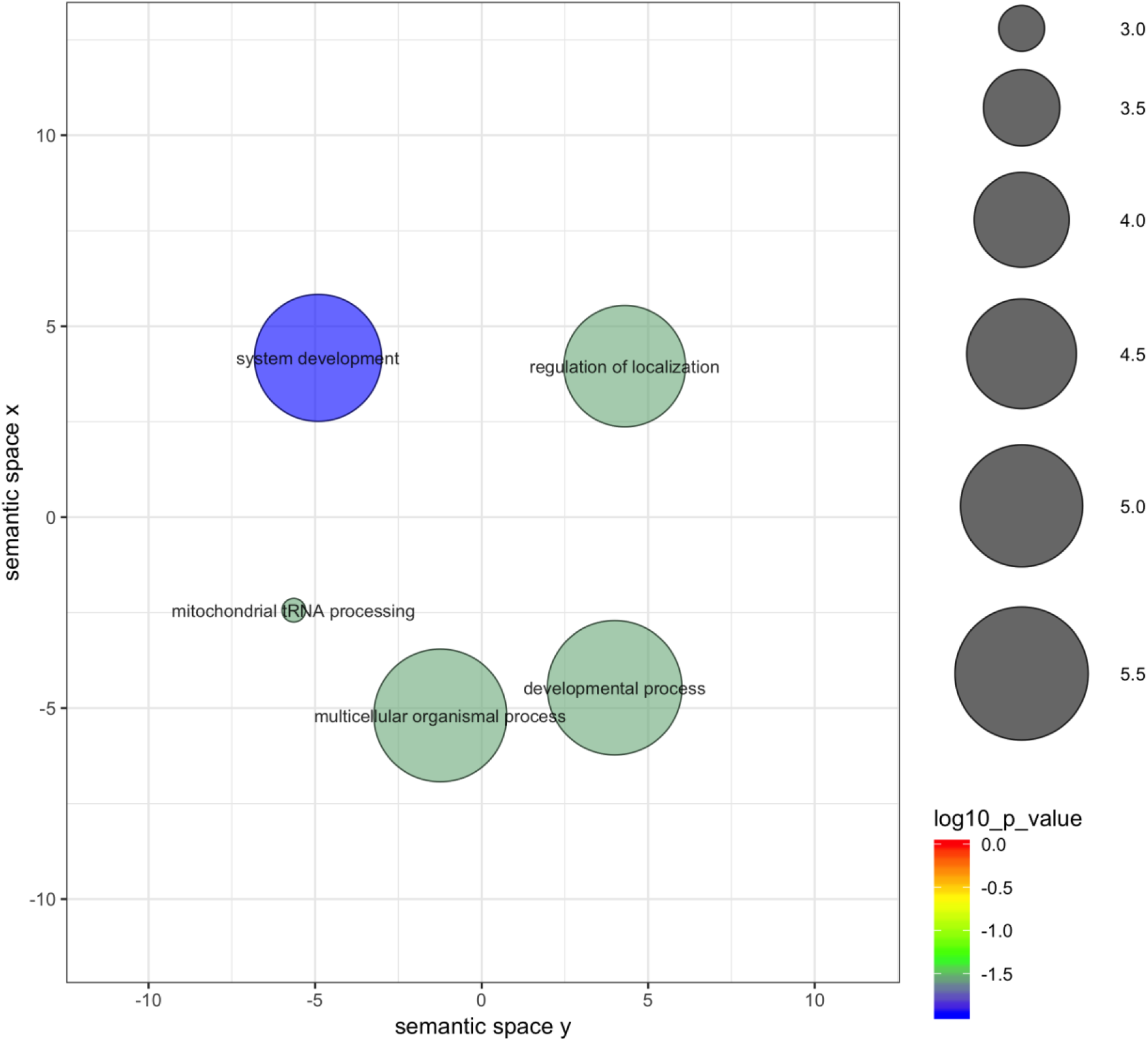
Results of the gene enrichment test for DMRs identified in the binomial regression in the CONTROL-HIGH comparison after merging semantically similar gene ontology categories.

**Figure 4.**
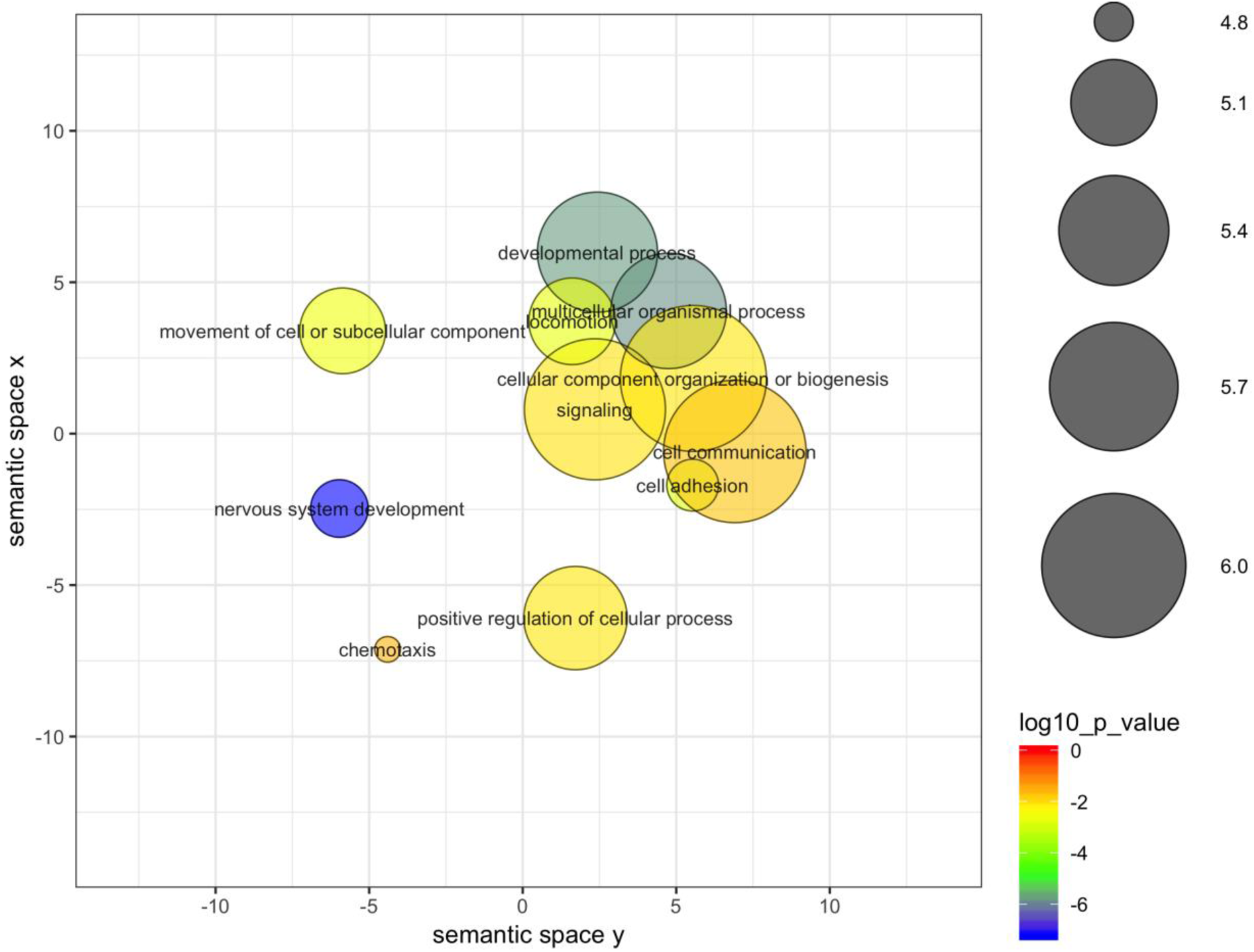
Results of the gene enrichment test for DMRs identified in the binomial regression in the CONTROL-SMELTER comparison after merging semantically similar gene ontology categories.

#### Differentially methylated CpG sites

We identified 96377 differentially methylated CpGs in the CONTROL-HIGH comparison and 129830 CpGs in CONTROL-SMELTER comparison using binomial GLM (p-level 0.05 Table 2). Almost equal proportion of the significant CpGs showed hypomethylation (45.9%) in CONTROL and hypermethylation (54.1%) in HIGH. Similarly, 52.5% of the significant CpGs were hypomethylated and 47.5% were hypermethylated CONTROL-SMELTER comparison (supplementary figure 5). 2789 (2.9%) of the significant CpGs showing hypomethylation, and 3022 (3.1%) showing hypermethylation were shared between CONTROL-HIGH and CONTROL-SMELTER, respectively (supplementary figure 5). Beta binomial GLM identified 16852 differentially methylated CpGs in CONTROL-HIGH comparison and 22669 CpGs in CONTROL-SMELTER comparison at p-level 0.05 (Table 2). For beta binomial regression the original p-values were recalculated based on the test statistics as implemented in the R-package fdrtool (Strimmer 2008b). Since the binomial regression method implemented in the R package Methylkit does not report test statistics the re-estimation of p-values were conducted only for the beta binomial regression. After re-estimation of p-values with fdrtool, 62789 and 67418 CpG sites (p<0.05) were differentially methylated in CONTROL-HIGH and CONTROL-SMELTER comparisons, respectively (Table 2). Of the significant CpGs, 28297 (45%) were hypomethylated and 34492 (55%) were hypermethylated in CONTROL-HIGH, and 36968 (55%) were hypomethylated and 30450 (45%) were hypermethylated in CONTROL-SMELTER. 946 (1.5%) of the significant CpGs showing hypomethylation, and 947 (1.5%) showing hypermethylation were shared between CONTROL-HIGH and CONTROL-SMELTER comparisons (Supplementary figure 5).

**Table 2.**
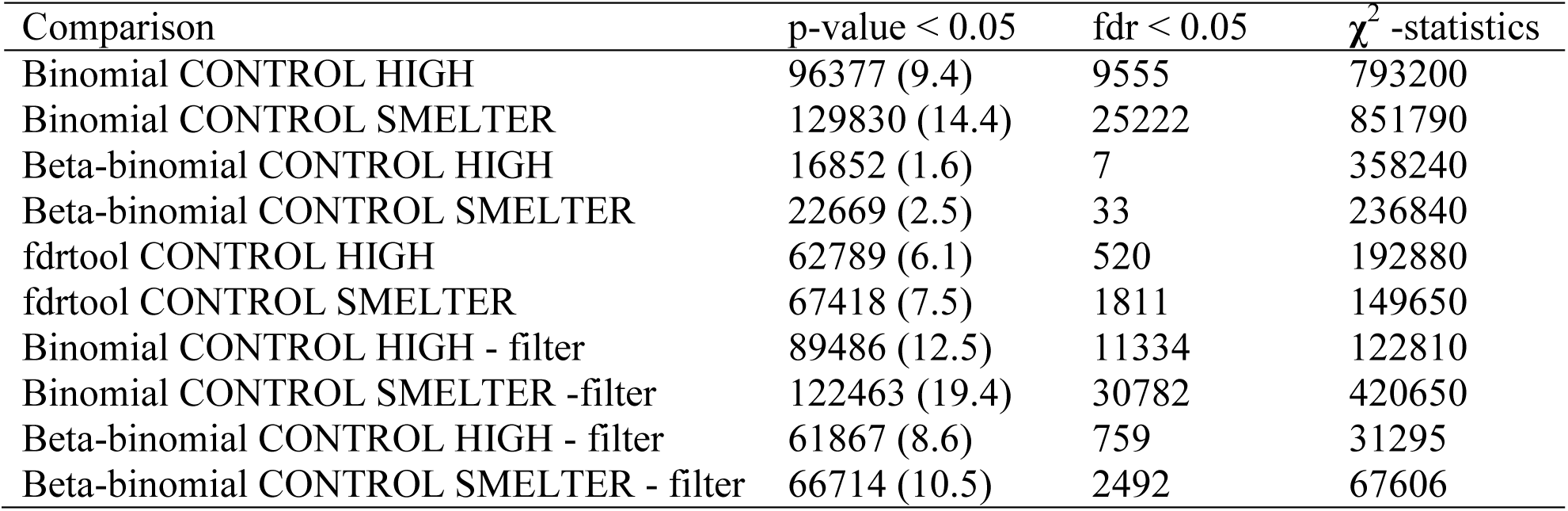
The number and percentage (in parentheses) of p-values less than 0.05 in different comparisons. Fdr refers to multiple testing correction using Benjamini & Hochberg (1995) method. The test statistics of goodness-of-fit test (**χ**^2^) of the p-value histograms indicated deviations uniform p-value distribution in unfiltered data, which improved with filtering (‘filter’).

The test statistics of goodness-of-fit test (Chi-square) of the p-value histograms indicated deviations from the uniform distribution in both methods (Table 2). However, the deviation in the beta binomial regression was smaller than the deviation in the binomial regression and the test statistics were lower when the p-values were re-calculated with fdrtool (Table 2). The deviations from the uniform distribution possibly indicate that our data do not fit to model assumptions or problems with the raw data (Strimmer 2008a, b). Also, methods for multiple testing assume uniform distribution (Strimmer 2008a). Therefore, we further investigated the deviation from uniform distribution by filtering the potentially uninformative CpGs as has been done previously on gene expression count data (Bourgon et al. 2010). More specifically, we calculated the mean of methylated counts (i.e. Cs) for each CpG across all individuals. We applied a threshold for the rank of mean methylated C counts and filtered out those CpGs that were causing the deviation from the uniform distribution (Supplementary figures 6 and 7) by keeping most of the significant CpGs. By removing 30% of the lowest C counts we recovered p-value distribution closer to the uniform distribution and a surplus for small (p<0.05) p-values (Figure 2, Table 1). The filtering was carried out using R package genefilter (Gentleman et al. 2018). The filtering approach also increased the number significant CpG sites after controlling for multiple testing (fdr <0.05) in all comparisons (Table 2).

#### Differentially methylated regions (DMRs)

Altogether, 336 DMRs were detected in binomial regression in CONTROL-HIGH comparison and 781 DMRs in CONTROL-SMELTER comparison 30 DMRs that had exactly the same starting position were shared between these two comparisons. 72 DMRs were detected in beta binomial regression in CONTROL-HIGH comparison and 159 DMRs in CONTROL-SMELTER comparison. Three DMRs were shared between these two comparisons. Of the DMRs identified in binomial regression, 54% and 40% of were hypomethylated in CONTROL compared to HIGH and SMELTER, respectively. In beta binomial regression 48% and 38% of the DMRs were hypomethylated in CONTROL compared to HIGH and SMELTER, respectively. This suggests that there was no clear pattern of hypo or hypermethylation in respect to pollution.

#### Annotation of the DMRs and CpGs

Altogether, the CONTROL-HIGH and CONTROL-SMELTER data sets were annotated to 13364 and 12972 genes, respectively. These data sets cover 72% and 70% of the total number of the annotated genes (18550) in Great tit genome. Of the 1023725 CpGs analyzed in the CONTROL-HIGH comparison, 683392 CpGs were found within genes (66.8%), 387735 (37.9%) in coding sequence and 637893 (62.3%) in exons. Of the 903449 CpGs analyzed in the CONTROL-SMELTER comparison 602997 CpGs were found within genes (66.7%), 334558 (37.0%) in coding sequence and 554421 (61.4%) in exons.

The DMRs identified in binomial regression in CONTROL-HIGH were annotated to 124 unique genes and CONTROL-SMELTER DMRs to 281 genes, excluding predicted genes. 46 of these annotated genes were shared between the comparisons. The DMRs from the beta binomial regression were annotated to 33 unique genes in CONTROL-HIGH and to 66 genes in CONTROL-SMELTER. Five of these genes were shared between the comparisons.

Gene enrichment analyses indicated 15 statistically significant (fdr < 0.05) gene ontologies in CONTROL-HIGH comparison (binomial regression) and 62 gene ontologies in CONTROL-SMELTER comparison. No statistically significant gene ontologies were found in either comparison among the DMRs identified in beta binomial regression. After merging semantically similar gene ontologies using REVIGO database, 5 and 11 enrichments remained in CONTROL-HIGH and CONTROL-SMELTER comparisons, respectively (Figure 3, Figure 4). Most of the gene ontologies were associated with developmental processes and were described under GO terms such as “system development” or “nervous system development” (Figure 3, Figure 4). Other categories involved cell-cell signaling or categories involving in transmitting information between cell and its surroundings (Figures 3, 4). Finally, we also report 10 DMRs with the largest differences in methylation levels (Supplementary Table 1). These included 12 genes (*POMC, ITGA11, LEKR1, USH2A, ZPR1, JMJD1C, ADAMTS3*, *PDE1C, TBP, PAPD4, GCC1* and *UTRN)* that may serve as potential candidates for further studies on the effects of pollution on organisms via DNA methylation.

## Discussion

We studied whether early-life exposure to pollution affects DNA methylation patterns in a wild great tit population. We found evidence that both direct lead exposure and experimental and anthropogenic pollution during post-hatching stage affect methylation levels of a small number (0.25-2.1%) genes from which we were able collect data, yet there was no consistent hypo or hypermethylation. The number of DMRs varied between binomial and beta binomial regression to a large extent such that binomial regression was more liberal than beta binomial regression. We found that genes associated with early developmental traits were overrepresented among the DMRs in binomial regression potentially linking methylation differences to biologically meaningful traits in birds living in polluted environments. Importantly, our results show that post-hatching environment modifies DNA methylation patterns in wild vertebrates.

### Direct effects of pollution on DNA methylation

Our data on fecal metal levels presented here, as well as data on bone lead levels (lead accumulates in bone) from the same broods (Eeva et al. 2014, Ruuskanen et al. 2015) shows that the HIGH group was indeed exposed to higher levels of lead than CONTROL during the post-hatching period. The measurements correspond to observed lead levels in polluted environments across Europe (Belskii et al. 1995a, Belskii et al. 1995b, Eeva and Lehikoinen 1996, Belskii et al. 2005, Berglund and Nyholm 2011), thus validating the effectiveness and environmental relevance of the lead exposure treatment.

The HIGH-CONTROL comparison represents direct effects of lead exposure post-hatching. There were no differences in the general methylation levels (hypo or hypermethylation) between the two groups, in contrast for example to previous epidemiological studies in humans (Pilsner et al. 2009, Wright et al. 2010). However, the identified GO terms that were found to be enriched using the binomial regression analysis suggest that high lead exposure affects methylation of genes associated with biological processes such as system development and developmental processes. In previous studies, similar developmental pathways have been identified in rodents, but also sex-specific differences reported (Singh et al. 2018). These results makes sense in the light of what is known from previous studies in the same study system. For example, in the HIGH lead treatment, vitamin A, retinol and stress protein levels were higher than in the CONTROL (Eeva et al. 2014, Ruiz et al. 2016). However, we expect that the patterns that we found in blood tissue are not only restricted to blood cells, but that the majority of the findings are likely to be similar for other tissues, as found previously in the study species (Derks et al. 2016, Verhulst et al. 2016). If the observed methylation differences lead to altered gene expression at the target genes, our results imply that the effects of pollution on such a variable set of genes may alter various developmental and cellular processes and ultimately health and phenotype.

### Indirect effects of pollution on DNA methylation

At the SMELTER site, birds were exposed to various pollutants, such as copper, nickel, cadmium and arsenic, originating from the nearby copper smelter (Eeva et al. 2014), with levels higher compared to CONTROL-HIGH. However, importantly, food quality and availability likely differ between CONTROL-SMELTER, as pollution reduces some important food sources such as caterpillars, and other insects in the area (Eeva and Lehikoinen 1996, Eeva et al. 2003). Furthermore, at the polluted site, individuals may be exposed to pollutants already pre-hatching as well as during post-hatching development. We found very little overlap (∼1-3%) in individual CpG sites between the CONTROL-HIGH and CONTROL-SMELTER comparisons. However, on the DMR level and their annotations showed some overlap indicating that the exposure to the lead treatment and to a larger collection of heavy metals at the smelter site can induce similar methylation changes. This suggests some direct effects of metals also at the smelter site. The gene ontology enrichments also mainly pointed that developmental processes were similar in these two comparisons suggesting that the overall effect of pollution is in the same direction. However, the majority of methylation differences in CONTROL and SMELTER are thus likely to be explained by by other elements than lead, and/or indirect effect of food, or their combination, either pre- or postnatally. Given that the metal concentrations observed at the SMELTER area are generally below the critical levels associated with sub-clinical effects (Berglund et al. 2012), indirect pollution effects via lower quality food may be likely (Eeva et al. 2005).

The number of DMRs between CONTROL and SMELTER were considerably higher than in CONTROL-HIGH comparison probably reflecting exposure to a more stressful environment, both nutritional stress and direct exposure to pollutants of various types. If the observed methylation differences lead to altered gene expression at the target genes (see below), they could contribute to the potential developmental problems associated with poor nutrition. For example, we found that SMELTER group showed lower growth rates, higher antioxidant enzyme and stress hormone levels, lower hematocrit and survival probability than CONTROL (Eeva et al. 2014, Rainio et al. 2015b, Ruiz et al. 2016).

The signal on differential methylation for genes related to nervous system development we detected in CONTROL-SMELTER comparison could potentially point to cognitive or behavioral changes. Parallel to our results, both prenatal lead and malnutrition have recently been found to influence methylation of genes in pathways associated with neuronal proliferation and differentiation in mice and embryonic cell models (Senut et al. 2012, Senut et al. 2014, Weng et al. 2014, Sanchez-Martin et al. 2015, Singh et al. 2018, Dou et al. 2019). In humans, captive animal models and wildlife, both early nutrition and metal exposure, particularly Pb, have well-documented detrimental effects on cognitive abilities and behavior that persist into adulthood (e.g., impaired learning, memory, increased aggression, hyperactivity Brown et al. 1971, Morgan et al. 2000, Burger and Gochfeld 2005, Carere et al. 2005, Arnold et al. 2007, Chen et al. 2012, Ruuskanen et al. 2015). Until now, the role of epigenetic mechanisms underlying such effects has not been thoroughly characterized. Our results can thus stimulate further research on the potential epigenetic mechanisms explaining the long-lasting influences of early-life adverse environment on behavioral and cognitive traits.

### Genes at differentially methylated regions

Among the set of 10 most differentially methylated regions across the treatment groups, we identified 12 genes (*POMC, ITGA11, LEKR1, USH2A, ZPR1, JMJD1C, ADAMTS3*, *PDE1C, TBP, PAPD4, GCC1* and *UTRN).* Of these 12, *POMC*, showed lower methylation (ca. 20%, thus theoretically higher expression) in methylation *both* in SMELTER and HIGH treatments compared to CONTROL. *POMC* (pro-opiomelanocortin) is a neuronal hormone, which is cleaved to multiple key by-products, including (i) corticotropin (ACHT), controlling the stress response, (ii) appetite control and (iii) b-endorphin (Marco et al. 2016). Methylation of *POMC* has been associated with nutritional state (in rats, Ramamoorthy et al. 2018), maternal under nutrition (in ovine Stevens et al. 2010) and offspring early-life stress (in mice, Wu et al. 2014). Here, we report for the first time that *POMC* methylation may mediate early-life stress (nutritional and/or metal exposure) also in a wild vertebrate population. Furthermore, methylation of another stress related gene, PDEC1 (phosphodiesterase 1C) was also decreased in SMELTER compared to control. Expression of PDEC1 gene has been found to be associated with aldosterone stress hormone in chicken (e.g. Fallahsharoudi et al. 2017).

The other differentially methylated genes in relation to metal exposure were related to (i) DNA damage: *JMJD1C* is a candidate histone demethylase and also plays a role in the pathway DNA-damage response (e.g. Watanabe et al. 2013). Our data suggests that its methylation was decreased (theoretical expression increased) in high lead exposure compared to control, which is logical given that lead exposure is likely to cause more oxidative stress and DNA damage (e.g. Wu et al. 2008, Rainio et al. 2015a). Other studies on Pb exposure also found differences in methylation in detoxification pathways (Sen et al. 2015); (ii) growth and development: *LEKR1* (Freathy et al. 2010), *ADAMTS3* (Janssen et al. 2016), *UTNR* (in mammals, e.g. Schofield et al. 1993). Furthermore, lead has specifically been found to impair neurodevelopment (e.g. Morgan et al. 2000, Burger and Gochfeld 2005). Our data shows that methylation of *ZPR1* (zinc finger protein gene), an important protein in neural development (e.g. Doran et al. 2006) was increased by ca. 20% (theoretical expression decreased) in lead exposure compared to controls, which may warrant further studies on *ZPR1,* lead exposure and neurodevelopment. Previous studies have reported alternation in methylation of other genes related to neurodevelopment, such as another zinc finger protein gene, *Zfp974* and *Zfp787, ARTN, C5aR1* (Dou et al. 2019), *Syt2, Prkg1, Pcdhb20, Slc2a3, Klhl1 and Snap29* (Singh et al. 2018) and *PAX1* and *MSI1* (Senut et al. 2014) (iii) transcription and intracellular processes: *TBP* is universal transcription factor required for all of the eukaryotic RNA polymerases (Shimada et al. 2003), *PAPD4* is a poly(A) RNA polymerase (Burroughs et al. 2010), while *GCC1* is associated with Golgi apparatus structure (Gosavi et al. 2018). Thus, these genes may serve as potential candidates for further studies on the effects of pollution on organisms via DNA methylation.

### Functional consequences of varying DNA methylation levels?

Importantly, when interpreting the potential functional consequences of the observed methylation differences, one needs to note that not all these genes with DMRs have been characterized in birds (and annotation has been done using mainly chicken and zebra finch gene models). Thus, the function of these genes is not well understood. Secondly, the link between DNA methylation and gene expression is not always straightforward (Jones 2012). However, we hypothesize that differential methylation at the observed sites affects gene activity and ultimately multiple cellular, developmental and physiological processes. Indirect evidence for a functional interpretation is provided by a recent great tit study using whole-genome bisulphite and RNAseq data. This study showed that across all genes, higher CG methylation at transcription start sites and within gene bodies was associated with lower gene expression (Laine et al. 2016). Finally, without detailed knowledge on gene function or differences in expression, it is difficult to judge whether the observed changes in methylation cause differences in phenotype and physiology we previously observed between HIGH lead exposure and CONTROL groups (Eeva et al. 2014, Rainio et al. 2015b, Ruiz et al. 2016). Therefore, follow-up studies are needed to investigate how the observed parameters are affected by differential methylation in one or more of the regions.

### Methodological considerations

We employed two commonly used methods to detect CpGs and evaluated their performance using p-value histograms. Either one of these methods did not recover uniform p-value histograms in our data when applied to overall coverage threshold of 10x, pointing out that the p-values are not reliable as such. When we applied a filtering approach, developed for gene expression count data, we were able to recover uniform distribution for both methods. Thus, it appears that uninformative counts i.e. low counts for methylated state can induce a clear deviation from the uniform distribution. In other words, the small methylation differences between treatment groups are potentially difficult to model using the two statistical approaches. While we applied an overall coverage threshold of 10x to our data, it seems that another filtering step is needed for C counts to recover uniform p-value distribution at least in our data set. Overall, the performance of the binomial and beta binomial regression reflects the outcome of previous studies on simulated and empirical data sets: binomial regression has been found to be more liberal in finding CpGs as compared to beta binomial regression (Dolzhenko and Smith 2014, Park and Wu 2016, Wreczycka et al. 2017). We found considerably more CpGs and DMRs in both comparisons using binomial than beta binomial regression. Overall, the comparison between the two methods is challenging in empirical data sets but both methods seem to recover uniform p-value distribution when uninformative CpGs are filtered out.

### Conclusions

In this study, we explored the environmental causes of epigenetic variation in an ecological, non-model organism, which is a novel and emerging research field. We found evidence that differentially methylated regions contain genes enriched for biologically meaningful processes and suggest potential targets for future research. Although we used a method that does not cover the whole genome, we were able to analyze methylation patterns covering most of the annotated genes in great tit genome. Thus, bisulfite sequencing (RRBS) can be a powerful and cost-efficient method in non-model species. However, the results were not consistent between binomial and beta binomial regression, which warrants caution when selecting analysis methods and interpreting results using different methods. Finally, the functional consequences of variable methylation patterns found in this study are yet to be discovered and a more comprehensive approach combining other molecular levels as well functional studies is needed.

## Acknowledgements

We thank Miia Rainio, Salla Koskinen, Tarja Pajari, Marjo Aikko, Orsolyia Palfi, Åsa Berglund and Jorma Nurmi for their efforts in helping us with field work. We also thank Finnish Centre for Scientific Computing (CSC) for providing computational resources, Center for Evolutionary Applications for molecular work and Finnish Functional Genomics Centre for sequencing services. Our study was financed by KONE foundation (SR) and Academy of Finland (TE: project 265859), Finnish Cultural Foundation (HM) and Turku University Foundation (SR and HM)

## Data accessibility

The sequence data are deposited in the SRA database under accession number PRJNA589705.

## Author contributions

SR, TE and HM designed the study. SR and TE collected the data. HM, SR and KvO designed the sequencing. HM conducted statistical and bioinformatic analyses. KvO and VL provided the genome resources. SR, HM, KvO and VL interpreted the data. All authors contributed to manuscript preparation.

---

**Supplementary figure 1.**
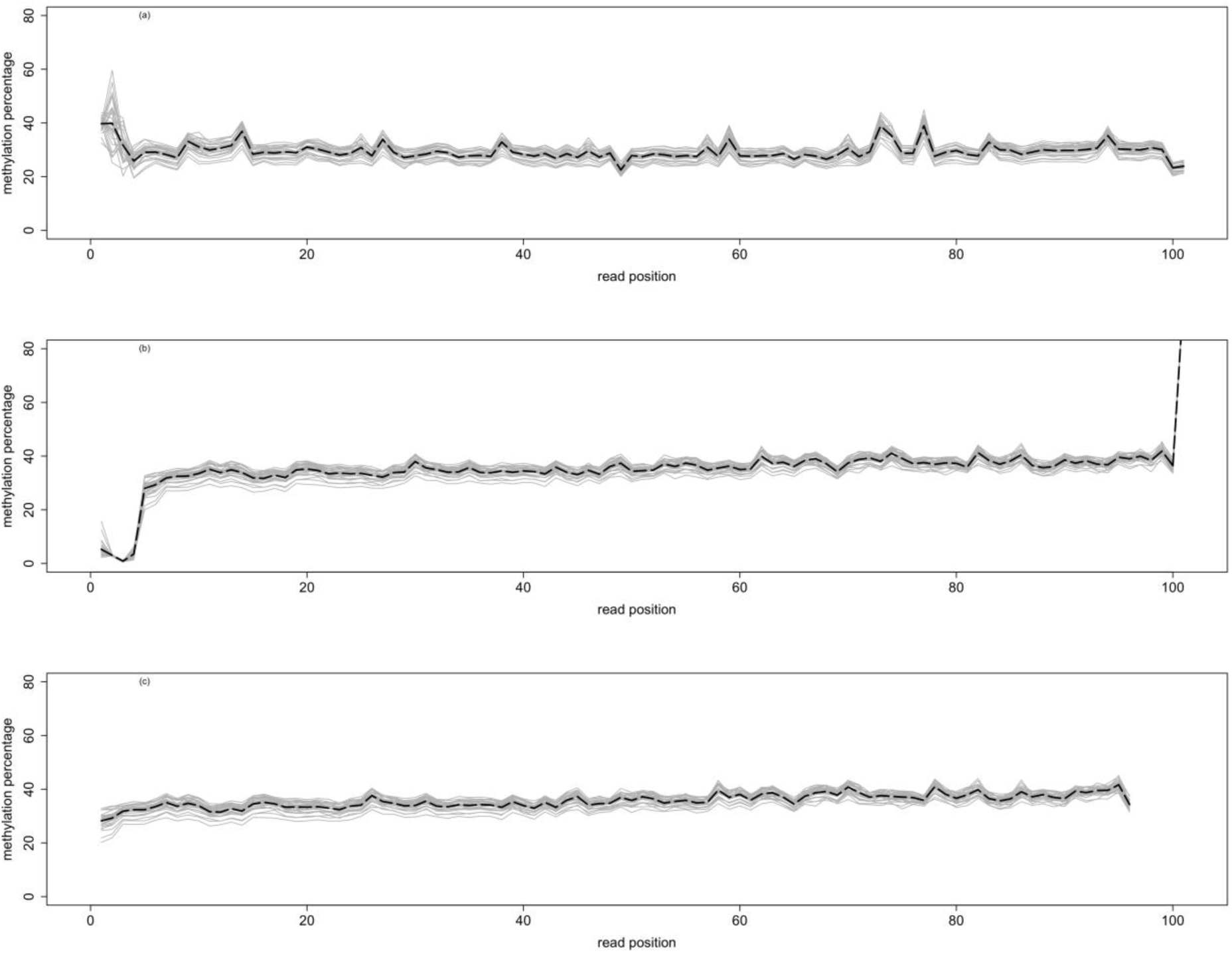
Methylation bias (M-bias) plots for read 1 (a) and for read 2 before (b) and after (c) cutting the bases showing lower or higher methylation than the other bases in the read. Four bases were cut from the beginning of the read 2 and one from the end of the read 2. The grey lines show the methylation percentage in all libraries along the position in the reads and the black line shows the mean methylation level across all libraries. On the x-axis is the position in the read and on the y-axis the methylation percentage.

**Supplementary figure 2.**
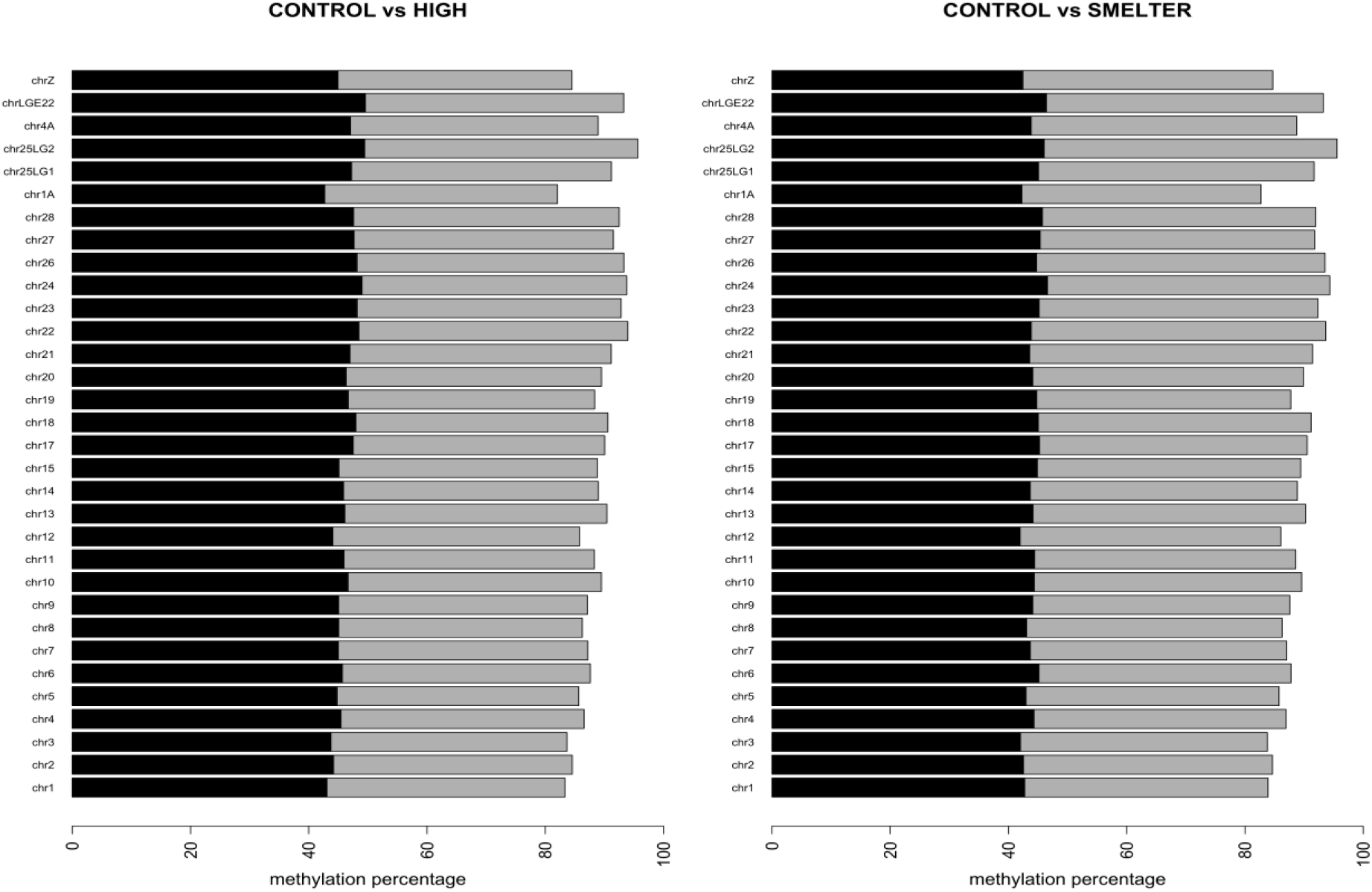
The percentage of hypo (black) and hypermethylated (grey) CpGs in Great tit major chromosomes. In (a) CONTROL and HIGH and in (b) CONTROL SMELTER comparisons.

**Supplementary figure 3.**
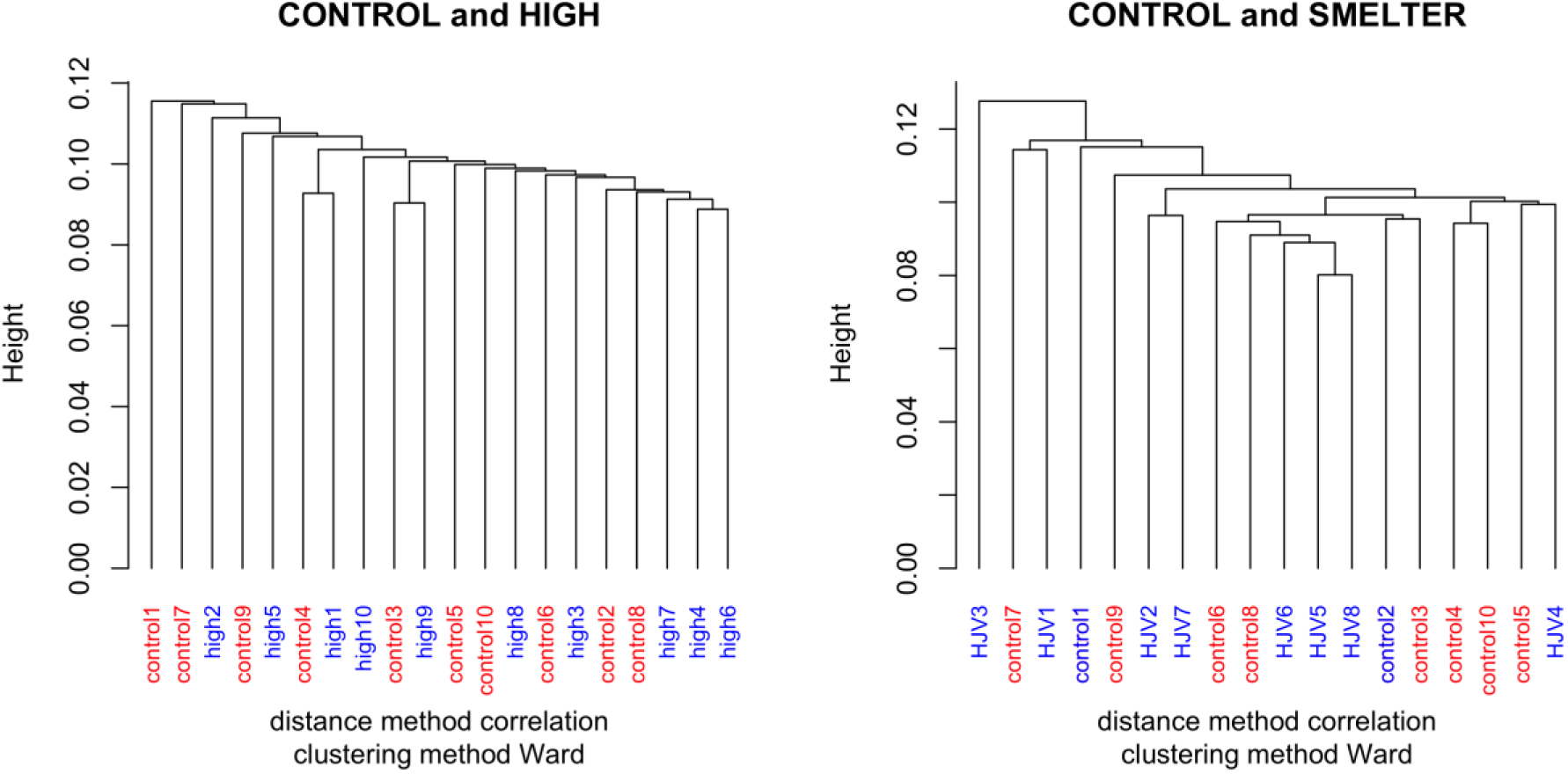
Hierarchical clustering of all individuals in (a) CONTROL HIGH and (b) CONTROL SMELTER comparisons.

**Supplementary figure 4.**
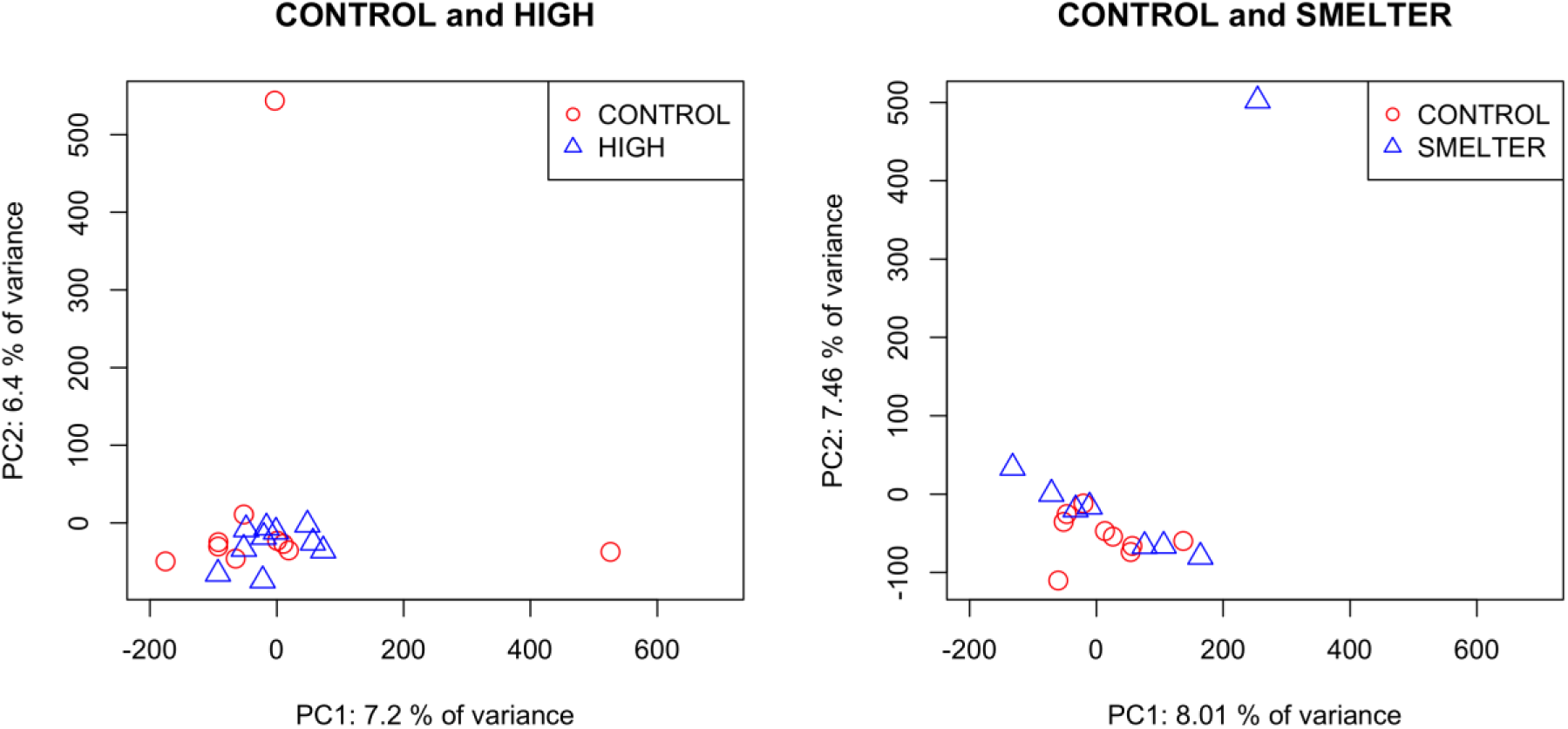
Principal component analysis of all individuals and CpGs in CONTROL HIGH (a) and in CONTROL SMELTER (b) comparisons. The x-axis shows the variance explained by PC1 and y-axis variance explained by PC2.

**Supplementary figure 5.**
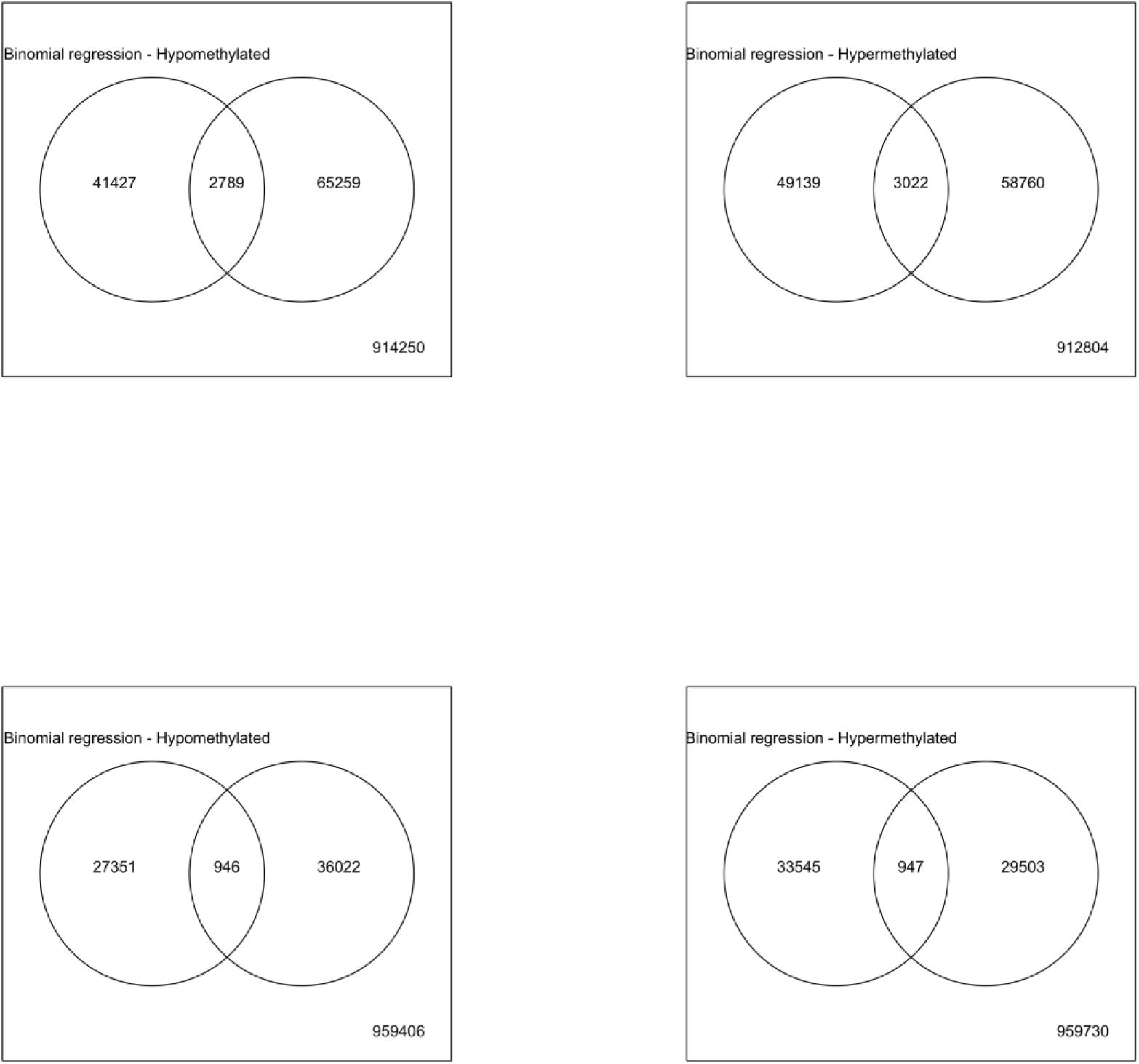
Venn diagrams showing the overlap of the significant CpGs between CONTROL-HIGH and CONTROL-SMELTER comparisons. In the upper panel is the overlap in binomial regression and in the lower panel the overlap in binomial regression.

**Supplementary figure 6.**
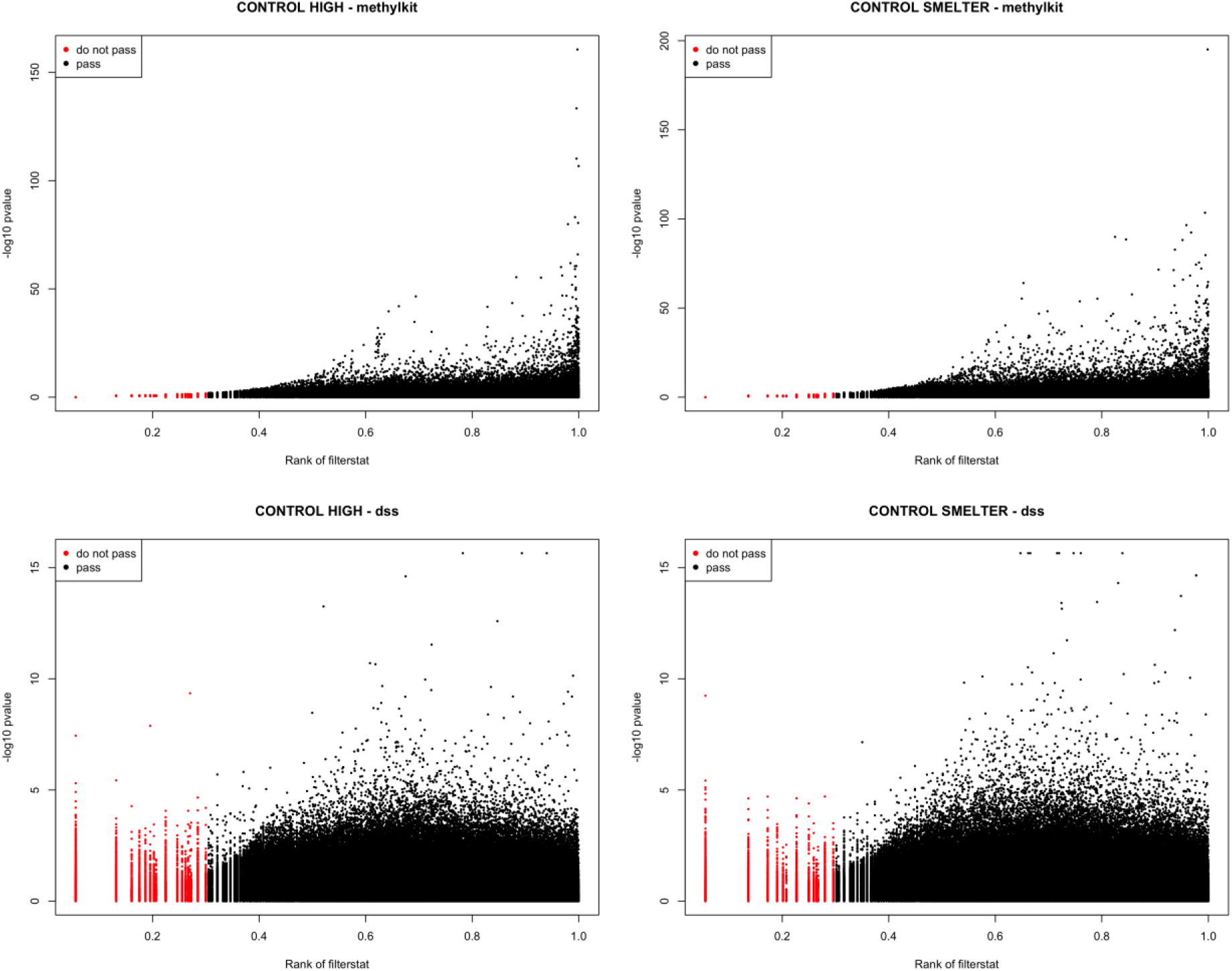
The principle of filtering uninformative CpGs. On the x-axis is the rank of filter statistics i.e. the mean coverage of Cs in each analyzed CpG. On the y-axis is the -log10 p-value obtained either from binomial regression (methylkit) or from beta binomial regression (dss). The red filled circles are CpGs not passing the filtering threshold (30%) while the filled black circles are CpGs passing the filtering threshold.

**Supplementary figure 7.**
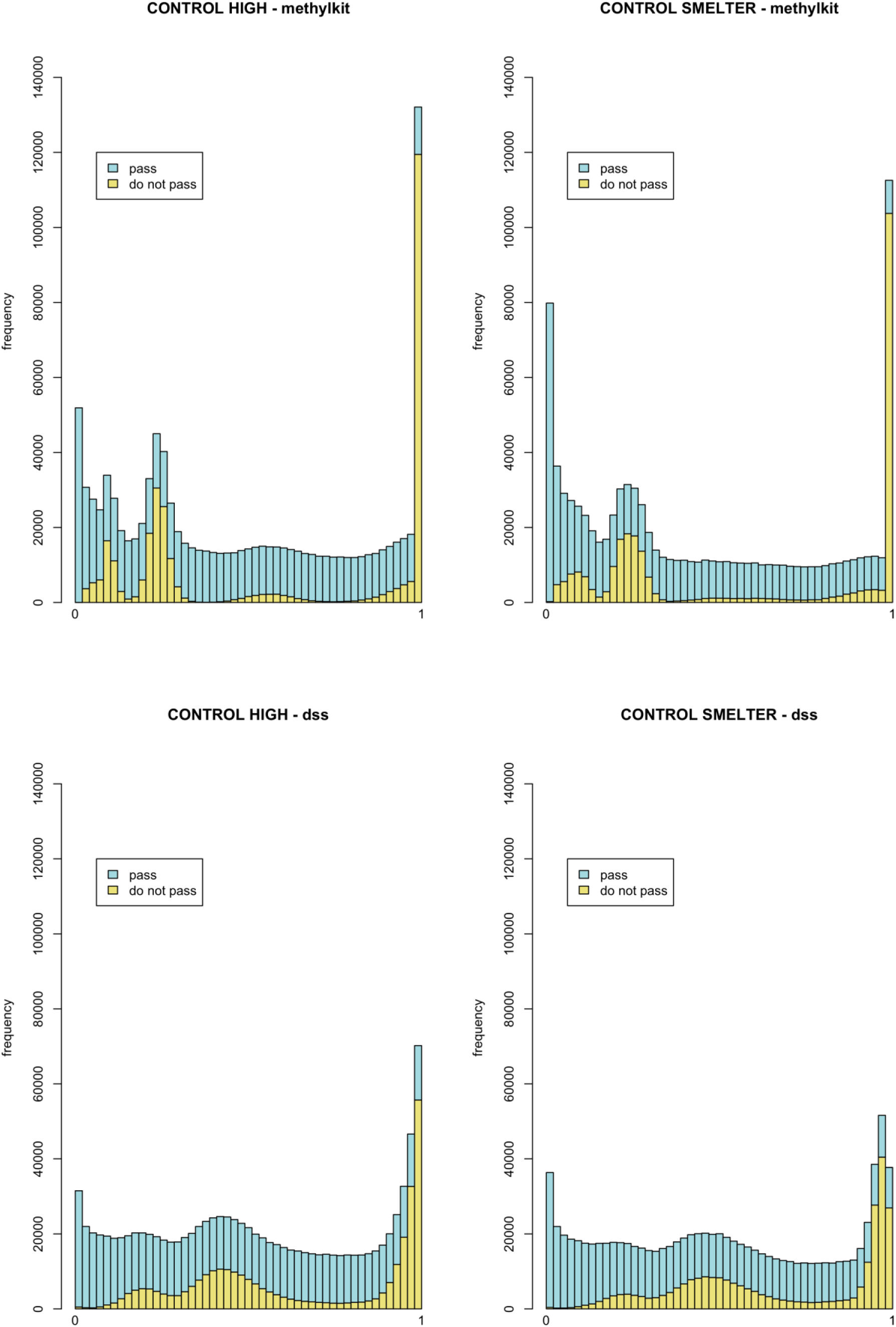
The filtering approach shown as p-value histogram. The yellow color shows the effect of removing low coverage CpG sites along the p-value distribution. The blue colour shows the p-values remaining after the low coverage filtering.

**Supplementary Table 1.**
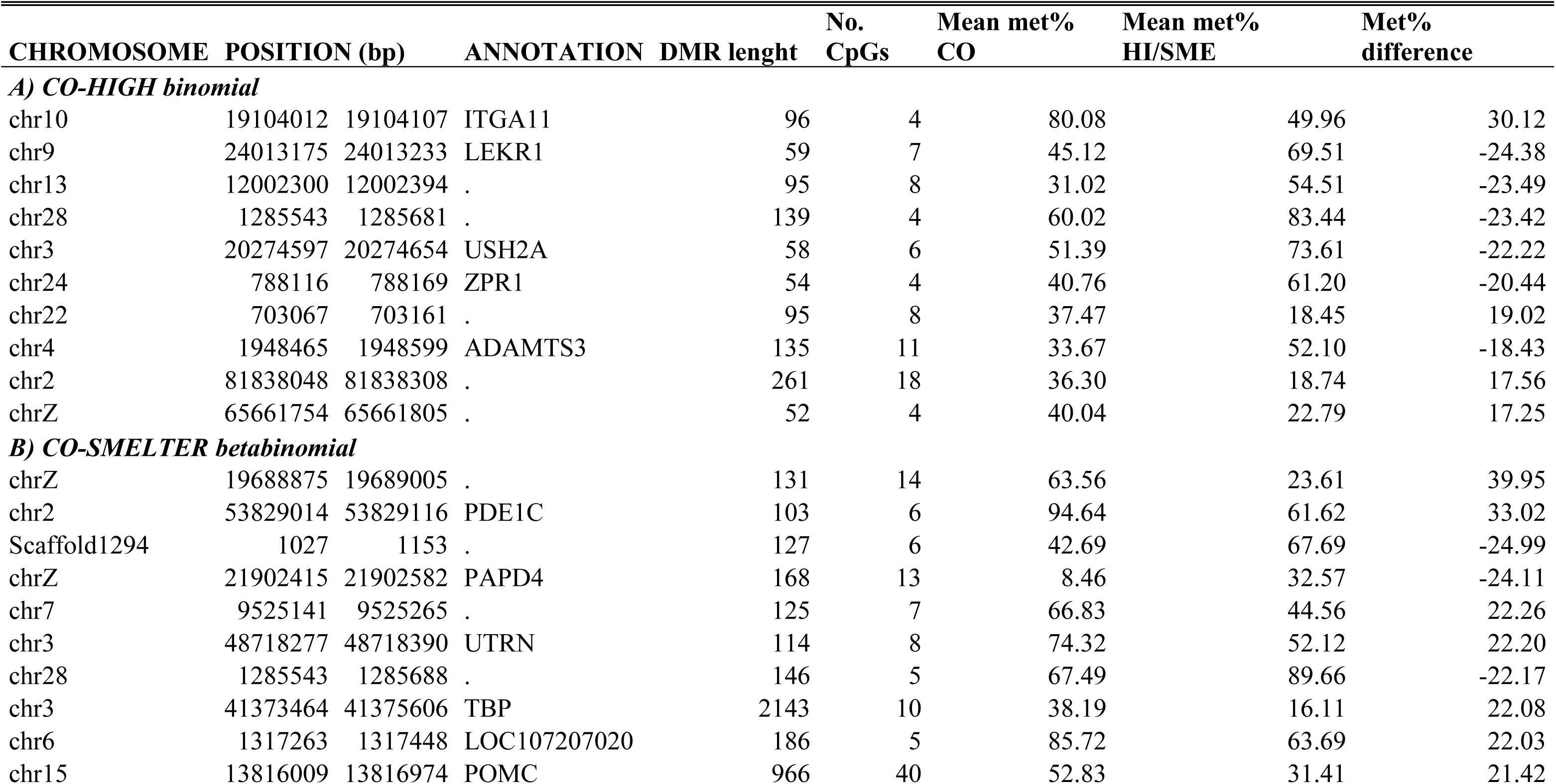
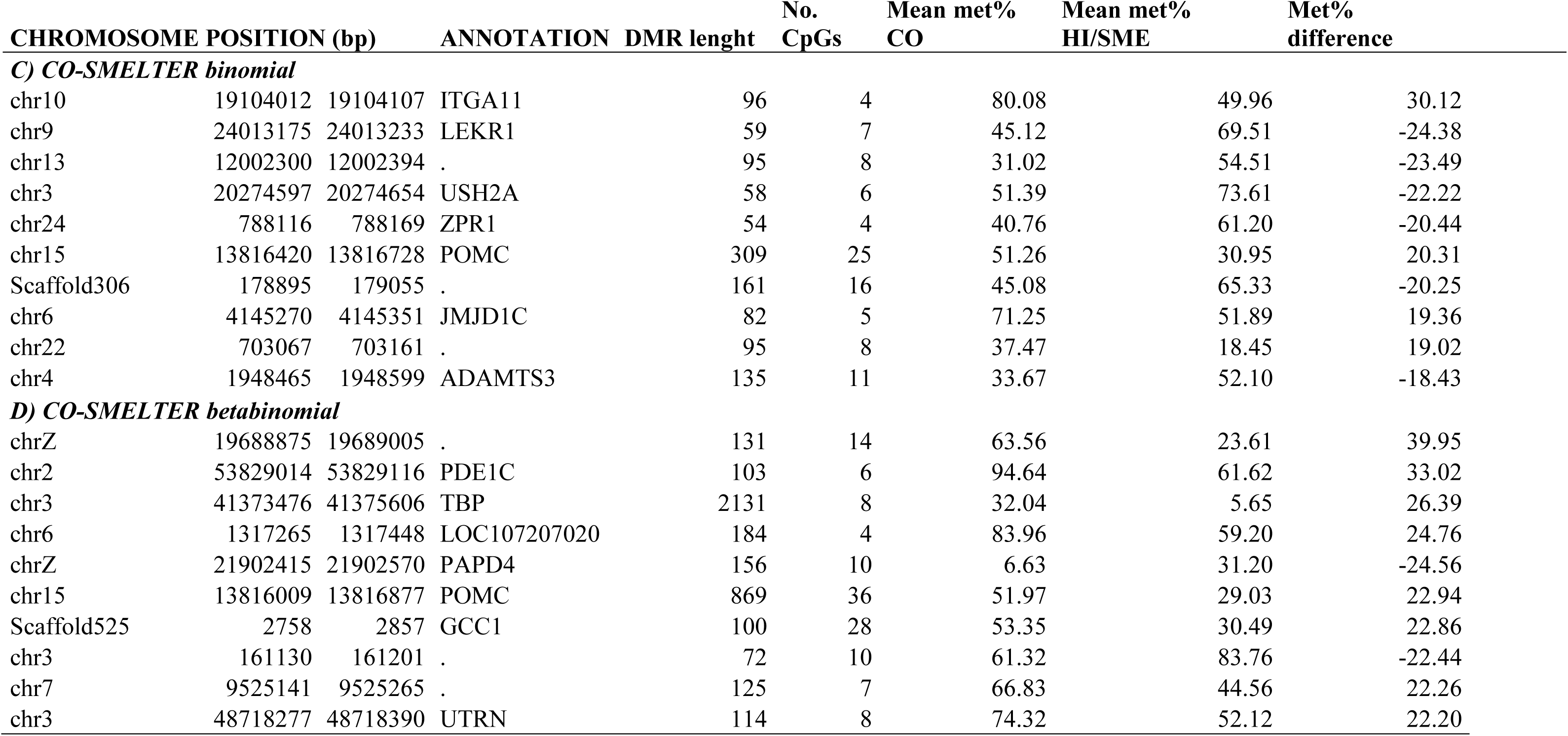
Top 10 DMRs for CONTROL-HIGH and CO-SMELTER comparisons, separately for binomial and betabinomial models.Mean Met% = average methylation percentage. Met% difference = difference, in percentages, in methylation between two groups. CO = control, HI = HIGH, SME = SMELTER

